# Y-Family DNA Polymerases Have Three Structural Elements That Promote Accurate dCTP Insertion And Minimize dATP/dGTP/dTTP Misinsertion Opposite a N^2^-dG Adduct of Benzo[*a*]pyrene

**DOI:** 10.1101/536987

**Authors:** Gabriel Sholder, Peter Tonzi, Sushil Chandani, Edward L. Loechler

**Affiliations:** Department of Biology, Boston University, Boston, MA 02215; New York University School of Medicine, New York, NY 10016; Novarus Discoveries Pvt. Ltd., Hyderabad 500073, India

## Abstract

To bypass DNA damage, cells have Y-Family DNA polymerases (DNAPs). One Y-Family-class includes DNAP κ and DNAP IV, which accurately insert dCTP opposite N^2^-dG adducts, including from the carcinogen benzo[*a*]pyrene (BP). Another class includes DNAP η and DNAP V, which insert accurately opposite UV-damage, but inaccurately opposite BP-N^2^-dG. To investigate structural differences between Y-Family-classes, Dpo4 (a canonical η/V-class-member) is modified to make it more κ/IV-like, as evaluated via primer-extension studies with a BP-N^2^-dG-containing template. Three protein structural elements are identified that promote fidelity. (1) Watson-Crick-like [dCTP:BP-N^2^-dG] pairing requires the BP-moiety to be in the minor groove. Thus, as expected, dCTP insertion is facilitated by having large openings in the protein surface that can accommodate BP-bulk in the minor groove. The BP-moiety is also in the minor groove during dATP and dTTP misinsertion, though evidence suggests that each of these three minor groove BP-conformations differ. (2) Plugging an opening on the major groove side of the protein suppresses dGTP misinsertion, implying BP-N^2^-dG bulk is in the major groove during Hoogsteen *syn*-adduct-dG:dGTP pairing. (3) Y-Family DNAPs have non-covalent bridges (NCBs) holding their little finger-domain in contact with their catalytic core-domain; dATP/dGTP/dTTP misinsertions are suppressed by the quantity and quality of NCBs, including one near and another distal to the active site on the minor groove side. In conclusion, three protein structural elements enhance dCTP and/or suppress dATP/dGTP/dTTP insertion; four different BP-adduct conformations are responsible for the four different dNTP insertional pathways opposite BP-N^2^-dG; generalizations about Y-Family structure are also considered.

## INTRODUCTION

Cells possess many DNA polymerases (DNAPs); e.g., humans, yeast (*S. cerevisae*) and *E. coli* have at least seventeen, eight and five, respectively.^*(1–3)*^ These DNAPs serve many functions; e.g., DNA damage caused by chemicals and radiation often block replicative DNAPs and to avoid such lethal blockage, cells possess lesion-bypass DNAPs, which conduct translesion DNA synthesis (representative recent references include: ^*(1, 2, 4–9)*^ with classic citations in reference 9). Most lesion-bypass DNAPs are in the Y-Family, which was the subject of a fine review.^*(7)*^ Human cells have three that are template-directed (hDNAPs η, ι and κ), while fission yeast has one (ScDNAPs η) and *E. coli* has two (EcDNAPs IV and V).

X-ray structures reveal that Y-Family DNAPs share a conserved catalytic region of ~350aa (representative references:^*(10–28)*^). All DNA polymerases resemble a right-hand with thumb, palm and fingers domains, which in Y-Family DNAPs make up the catalytic core (CC-Domain).^*(7)*^ Y-Family DNAPs grip DNA with another domain, ^*(10–12)*^ often called the “little finger” (LF-Domain). Y-Family DNAPs have “stubby” fingers, leaving more solvent accessible surface around the template/dNTP-binding pocket,^*(4)*^ which can accommodate adduct/lesion bulk. A representative X-ray structure of Dpo4 from *Sulfolobus solfataricus* (the best-studied Y-Family DNAP) is shown in Figure 1A, as viewed from the minor groove with its CC-Domain in green and LF-Domain in yellow. Representative structures are also shown for SaDbh (Figure 1B), hDNAP κ (Figure 1C) and EcDNAP IV (Figure 1D).

**FIGURE 1:**
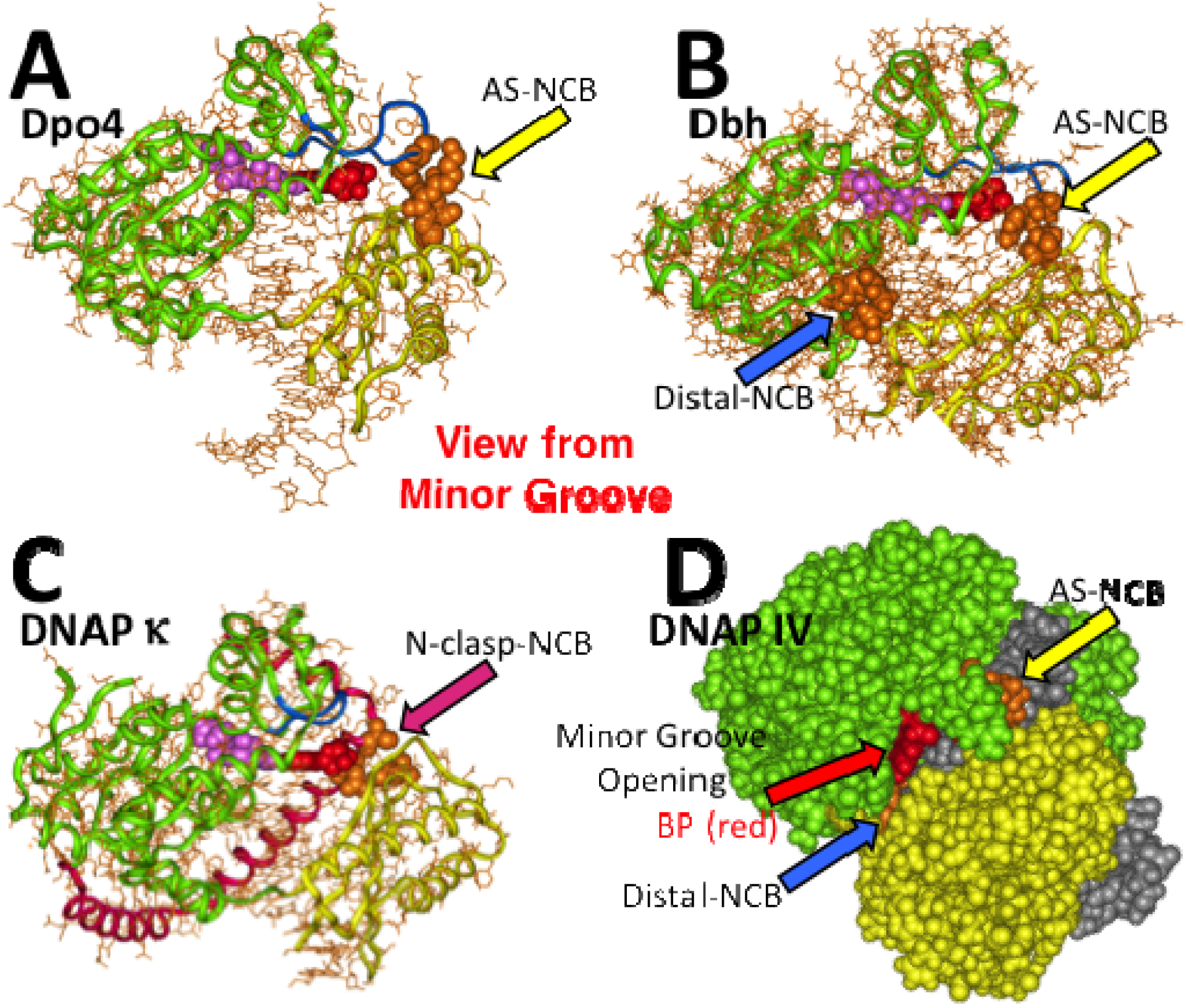
Views from the minor groove side of the active site for the Y-Family DNAPs: SsDpo4 (Panel A), SaDbh (Panel B), hDNAP κ (Panel C) and EcENAP IV (Panel D). The CC-Domains are shown in green and the LF-Domains are shown in yellow. Amino acids involved in the formation of noncovalent bridges (NCBs) between the CC-Domain and the LF-Domain are shown in brown, which can be of three types: near the active site (AS-NCB, yellow arrow) and distal to the active site on the minor groove side (Distal-NCB, blue arrow), or on the major groove side near the active site (N-clasp-NCB, scarlet arrow), as is present uniquely in hDNAP κ, because of its N-clasp (scarlet structure). X-ray sources of structures include SsDpo4 ^*(24)*^ and hDNAP κ, ^*(13)*^ while modeled structures are used for SaDbh and DNAP IV ^*(72)*^ for the following reasons. The X-ray structures for SaDbh ^*(27, 73, 76)*^ are with undamaged DNA containing a one-nucleotide bulge, which prevents AS-NCB formation; the structure shown is modeled with the DNA bulge removed, which allowed the AS-NCB to reform during molecular dynamics. Though X-ray structures exist for DNAP IV ^*(17, 26)*^, they do not have a BP-adduct, so we include our model with –BP-dG (red),^*(72)*^ which can be seen through the opening on the minor groove side of the active site. DNA is gray.

Steps in the mechanism of Y-Family DNAPs have been proposed for both protein structural changes based on a series of X-ray structures^*(29, 30)*^ and for chemical catalysis based on theoretical studies.^*(31)*^ Steps in Y-Family DNAP bypass of several lesions have been identified based on kinetic analysis,^*(32, 33)*^ hydrogen-deuterium exchange mass spectrometry,^*(34, 35)*^ and single molecule FRET.^*(36)*^

Our work has focused on benzo[*a*]pyrene (BP), ^*(37–41)*^ which is a well-studied DNA damaging agent that is a potent mutagen and carcinogen, and an example of a polycyclic aromatic hydrocarbon (PAH), a class of ubiquitous environmental substances produced by incomplete combustion.^*(42)*^ PAHs in general and BP in particular induce the kinds of mutations that may be important in human cancer (^*(43)*^ and references therein). Figure 2 shows the major BP DNA adduct (+BP-dG) and its mirror image (–BP-dG), both of which form at N^2^-dG.

**FIGURE 2:**
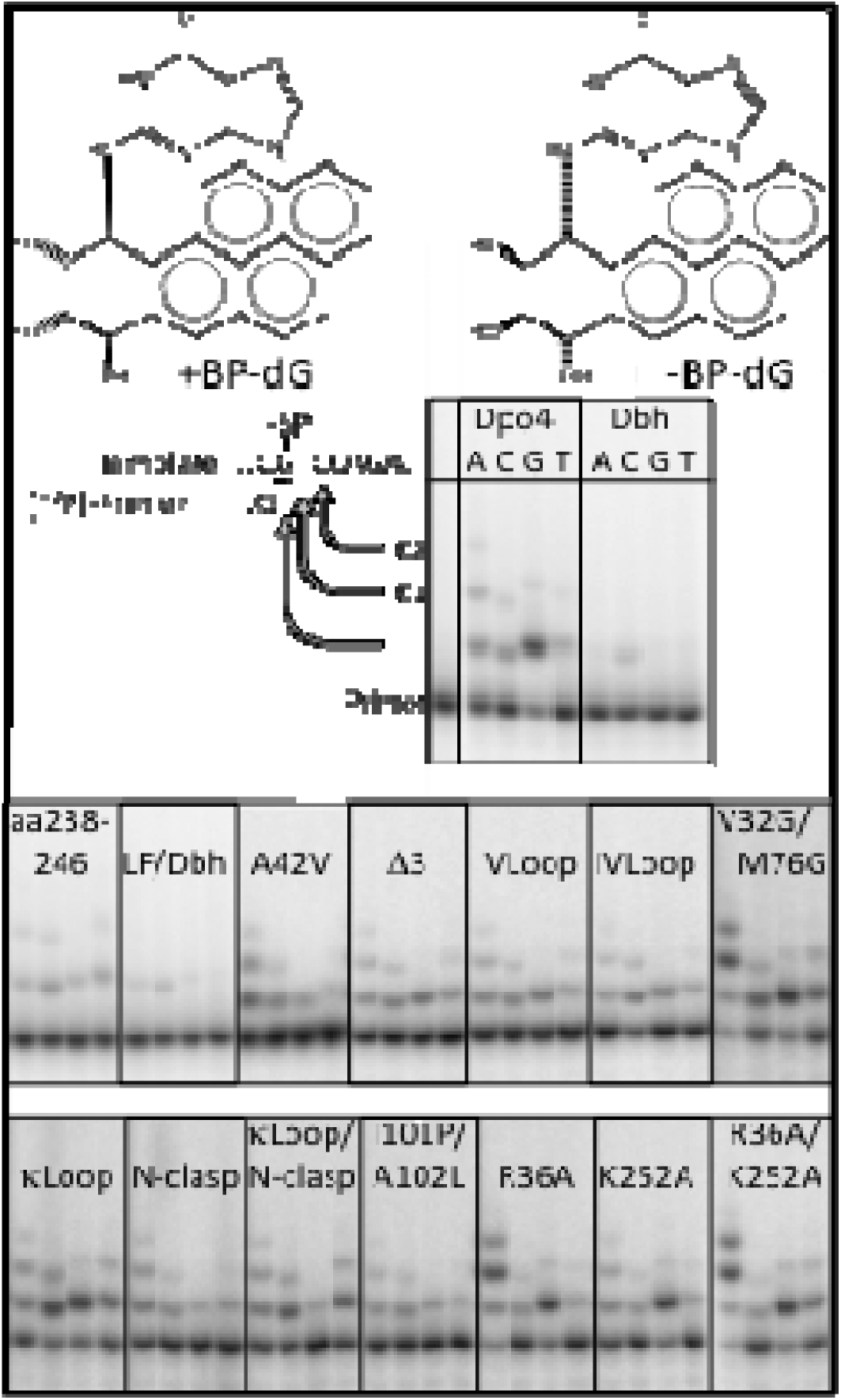
Structures of +BP-dG and –BP-dG, along with a portion of the primer/template sequence and a phophorimage showing Y-Family DNAPs SsDpo4, SaDbh and derivatives incorporating dATP, dCTP, dGTP and dTTP incorporation into a ^32^P-containing primer. Primer alone is shown in lane 1. Images for the SsDpo4 set, the SaDbh set and the primer were in the same gel (same images as in reference 37), but were repositioned in the figure. In addition, typical phosphorimages for each of the DNAPs listed in Table 2 are shown. The images come from many different experiments. Four lanes are shown for each DNAP arranged (left-to-right) dATP, dCTP, dGTP and dTTP, with incubations for 60min, 60min, 20min and 60min, respectively, except R36A/K252A-Dpo4, whose dGTP reaction was 5min.

Based on amino acid sequence, Y-Family DNAPs are divided into six major groups. ^*(7)*^ Based on insertional patterns opposite many lesions, classes seem to exist.^*(44)*^ The κ/IV-class correctly inserts dCTP opposite N^2^-dG adducts, including from +BP-dG and –BP-dG, ^*(28, 37, 45–56)*^ as well as from N^2^-dG adducts from endogenous sources. ^*(57, 58)*^ IV/κ-class-members also correctly insert opposite other lesions. ^*(59–62)*^ κ/IV-class-members do not readily insert opposite UV-damage and tend to be relatively accurate in the replication of undamaged DNA. ^*(63)*^ κ/IV-class-members include hDNAP κ, EcDNAP IV and SaDbh. η/IV-class-members are relatively inaccurate in replicating undamaged DNA and insertion opposite N^2^-dG adducts, but are accurate with UV-damage, with thymine dimers being the best studies example. ^*(44)*^ η/V-class-members include hDNAP η, UmuC (polymerase subunit of EcDNAP V) and SsDpo4.

**Table 2:** Pseudo First-Order Rate Constants for dNTP Insertion Opposite –BP-dG for the DNA Polymerases wt-Dpo4 and wt-Dbh, as well as Derivatives of Dpo4.^1,2^

| Wild Type | dATP |  | dCTP |  | dGTP |  | dTTP |  |
| --- | --- | --- | --- | --- | --- | --- | --- | --- |
| wt-Dpo4 | 11.2 | 0.25 | 5.1 | 0.25 | 73.8 | 0.20 | 3.8 | 0.15 |
| wt-Dbh | 0.64 | 0.029 | 2.0 | 0.10 | 1.1 | 0.29 | 0.49 | 0.29 |
| Dpo4 Derivatives | Major Groove Considerations |  |  |  |  |  |  |  |
| A42V | 12.0 | 0.10 | 4.9 | 0.081 | 7.4 | 0.21 | 3.4 | 0.24 |
| Δ3 | 9.7 | 0.27 | 5.5 | 0.23 | 4.8 | 0.41 | 3.5 | 0.29 |
| IVloop | 4.9 | 0.19 | 9.6 | 0.21 | 6.7 | 0.21 | 2.7 | 0.31 |
| Vloop | 9.8 | 0.16 | 5.2 | 0.075 | 7.8 | 0.12 | 3.0 | 0.23 |
| N-Clasp | 12.2 | 0.21 | 7.2 | 0.24 | 8.1 | 0.12 | 3.8 | 0.29 |
|  | Minor Groove Opening Size Considerations |  |  |  |  |  |  |  |
| V32G/M76G | 39.4 | 0.20 | 12.3 | 0.13 | 55.1 | 0.12 | 11.3 | 0.12 |
| κloop | 11.0 | 0.20 | 21.8 | 0.19 | 70.9 | 0.26 | 13.0 | 0.11 |
| N-Clasp/κloop | 13.5 | 0.15 | 13.1 | 0.071 | 7.6 | 0.11 | 12.0 | 0.036 |
|  | AS-NCB Considerations |  |  |  |  |  |  |  |
| R36A | 36.6 | 0.19 | 6.7 | 0.11 | 71.0 | 0.25 | 4.2 | 0.17 |
| K252A | 9.8 | 0.049 | 5.0 | 0.16 | 67.7 | 0.12 | 4.3 | 0.21 |
| R36A/K252A | 50.3 | 0.38 | 6.5 | 0.21 | 276. | 0.28 | 10.3 | 0.57 |
|  | Distal-NCB Considerations |  |  |  |  |  |  |  |
| LF/Dbh | 1.4 | 0.15 | 2.1 | 0.12 | 2.2 | 0.063 | 0.54 | 0.35 |
| aa238-246/Dbh | 5.2 | 0.24 | 5.2 | 0.29 | 10.3 | 0.38 | 2.0 | 0.21 |
| I101P/A102L | 4.0 | 0.36 | 4.8 | 0.94 | 21.2 | 0.29 | 3.1 | 0.80 |

<sup>1</sup> Pseudo first-order rate constants are reported for dNTP insertion opposite a –BP-dG adduct in a primer/template (left-most column for each dNTP) using standard conditions (dNTPs/100μM and DNAP/124nM, at pH 7.6 and 37°C); e.g., $11.2 \times 10^{-3} \text{ min}^{-1}$ is the value for Dpo4 with dATP. All values are the average of at least three determinations and are reported [ $\times 10^{-3} \text{ min}^{-1}$ ]. Purified proteins included wt-Dpo4, wt-Dbh and Dpo4 mutant derivatives (Table 1). Methodological details are summarized in Supplemental Materials and described in reference <sup>(40)</sup>.
<sup>2</sup> Standard deviations are reported in the right-most columns for each dNTP and are expressed as a fraction of the rate constants; e.g., for wt-Dpo4 with dATP, the true standard deviation ( $2.8 \times 10^{-3} \text{ min}^{-1}$ ) is divided by its pseudo first-order rate constant ( $11.2 \times 10^{-3} \text{ min}^{-1}$ ) to give its fractional standard deviation (i.e., 0.25).

Insertional preference differences for κ/IV-class-members versus η/V-class-members must reflect an interplay between protein structural differences and adduct structural/conformational differences. Five X-ray structures exist for several BP-adducts in Dpo4, ^*(23, 24)*^ and each structure is different. The adduct-dG can be either in the *anti*-orientation or *syn*-orientation, which places the BP-pyrene moiety in the minor or major groove, respectively. The BP-pyrene moiety can also be intercalated. Furthermore, the adduct-dG can be paired with a base in the opposite strand or be extra-helical. Finally, only one structure has the adduct dG-moiety interacting with an active site dNTP, while the others have the [n+1] base pair in the active site. These Dpo4 structures would most likely relate to mutagenic pathways. X-ray structures in hDNAP κ show a likely structure for correct dCTP insertion opposite +BP-dG, which has more-or-less normal Watson-Crick pairing with the +BP-moiety in the minor groove.^*(28)*^

Thus, X-ray studies reveal a diverse set of structural possibilities for BP-dG adducts, which provide tantalizing mechanistic possibilities, although this complexity makes the task of associating a particular structure/conformation with a particular dNTP insertion pathway daunting. To provide additional clues about the relationship between structure and mechanism, X-ray studies can be complemented by structure/activity studies, which can provide insights about transition state structures that control rates of reaction and, thus, mechanism.^*(64)*^ We take a structure/activity approach herein to probe dNTP insertion mechanism opposite –BP-dG, which we used in two prior studies.^*(40, 65)*^

Regarding misinsertion mechanism, dNTP insertion by Dpo4 has been studied opposite +BP-dG.^*(23)*^ dATP, dGTP and dTTP misinsertion dominated in 5′-TG, 5′CG and 5′-AG sequences respectively, which is consistent with a mutagenic insertion mechanism being dictated by the 5′-neighboring base via a slippage-type process.^*(41)*^ However, studies with +BP-dG and other BP-adducts have shown that mutagenic outcome being defined by the 5′-base is the exception rather than the rule (reviewed I, reference 41). In addition, we recently showed that—though Dpo4 does misinsert dATP opposite –BP-dG in a 5′-TG sequence context—dGTP is faster (~6.6-fold), while dTTP misinsertion, as well as correct dCTP insertion, are also significant. ^*(40, 65)*^ Thus, on balance, the nature of the 5′-base does not principally define insertion.

Another key question is: How does a single adduct direct multiple insertional events? For example, (1) does Dpo4 incorporate dATP, dCTP, dGTP and dTTP, ^*(40)*^ because it is unable to interpret the –BP-dG adduct, such that it merely incorporates whatever dNTP happens to be in its active site, or (2) are there mechanistically distinct incorporation pathways?

We have been investigating such mechanistic questions, principally by making structural modifications in wt-Dpo4, which is in the η/V-class and has low fidelity during BP-N^2^-dG adduct bypass,^*(23, 40, 65)*^ to generate Dpo4-derivatives that behave more like κ/IV-class-members, with increased rates (i.e., “activity”) of dCTP insertion and decreased rates of dATP, dGTP and dTTP misinsertion opposite –BP-dG. In one study with wt-Dpo4, several Dpo4 mutants and wt-Dbh,^*(40)*^) reactions were shown to follow simple Michaelis-Menten kinetics with respect to [dNTP]. Subsequently, the structural basis for mechanistic differences between Dpo4 and DNAP IV was investigated by studying the kinetics of various chimeric and mutant derivatives.^*(65)*^

The study herein extends this approach by modifying structural elements in Dpo4 to make it more like Dbh, DNAP IV and DNAP κ. We find the following. (1) The –BP-moiety is in the major groove during dGTP misincorporation. In contrast the –BP-moiety is in the minor groove during dCTP, dATP and dTTP misincorporation, although evidence suggests that each of these adduct conformations is different. Thus, findings are most consistent with four different adduct –BP-dG conformations being responsible for each of the four reactions involving dCTP, dATP, dGTP and dTTP. (2) The size of an opening on the minor groove side of the active site influences the rate of correct dCTP insertion opposite a BP-dG-adduct. This is sensible because the BP-moiety must be in the minor groove to form a more-or-less normal adduct-dG:dCTP Watson-Crick-like base pair, as first predicted by molecular modeling,^*(66)*^, and as observed in X-ray structures.^*(28)*^ (3) dATP, dGTP and dTTP misinsertions are suppressed by non-covalent bridges (NCBs) that hold the LF-Domain in contact with the CC-Domain, including one near the active site and a second that is distal to the active site. (4) Structural features are identified that κ/IV-class-members use to suppress misinsertions opposite adducts having bulk in the major groove. (5) Some of the Y-Family structural features described herein appear to be present in other Y-Family DNAPs. (6) The conclusions in this study are fundamentally consistent with those made in our previous studies.^*(40, 65)*^

## EXPERIMENTAL PROCEDURES

Vectors containing genes of interest have been described, along with methods to generate derivatives.^*(40)*^ The one exception is N-clasp-Dpo4, which contains the first 100aa of human DNAP κ fused to Dpo4. The vector pBJ733 ^*(67)*^ was PCR amplified with primers to give an appropriate N-clasp-containing fragment flanked by NdeI sites at each end. The fragment and wt-Dpo4-pET15B were each digested with NdeI, gel purified and ligated together. This method introduced an extra His residue between S100 of the N-clasp and M1 of Dpo4, which was removed via site-directed mutagenesis. Amino acid sequences of all derivatives are given in Table 1. Except as noted below, all other experimental procedures were described previously,

**Table 1:**
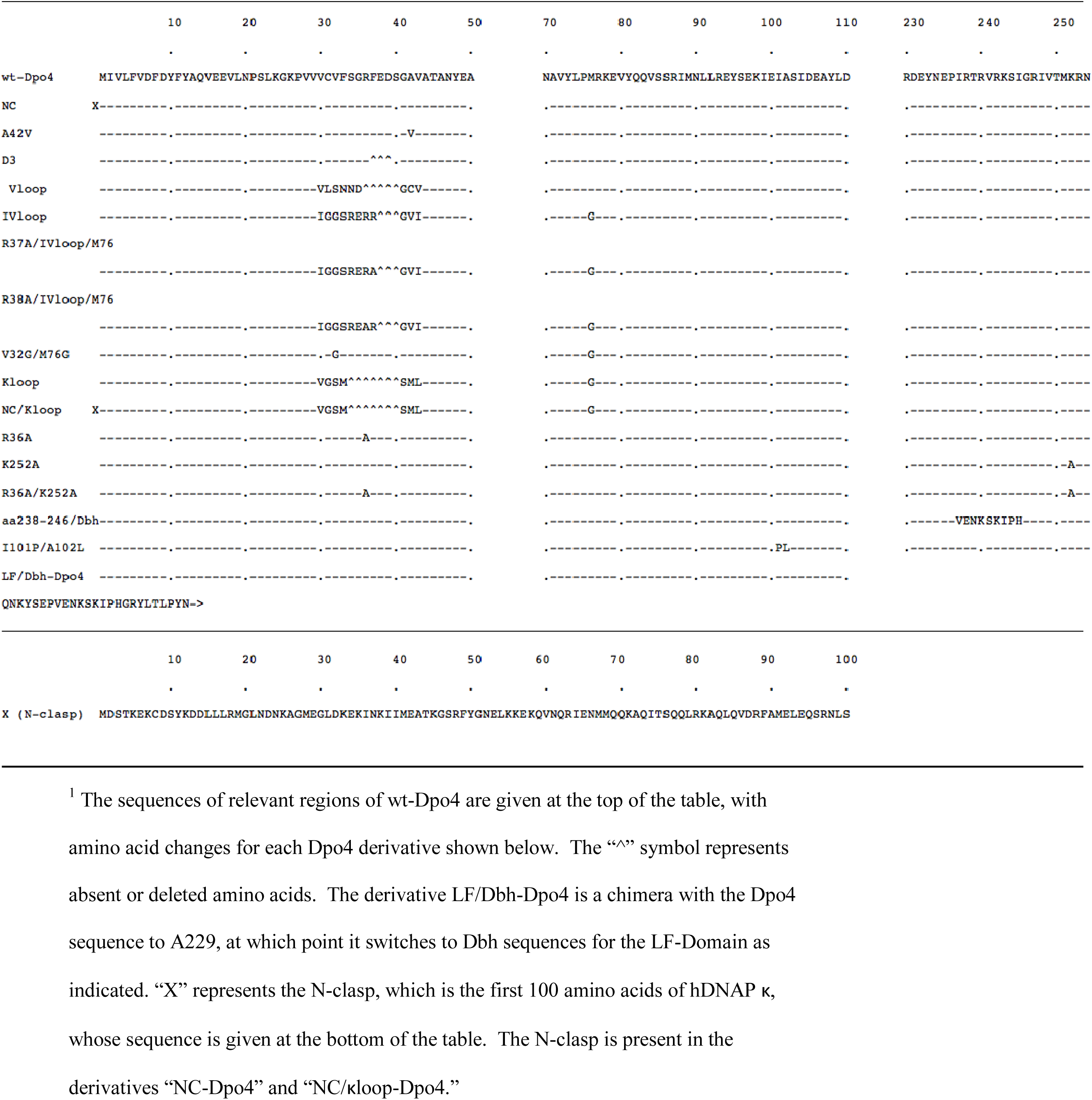
Sequences of Dpo4 Derivatives Used in This Work.^1^

^*(40)*^ including the method to accurately determine rate constants, which is summarized in Supplementary Materials. Primer-extension assays were at pH 7.6 and 37°C. To obtain measurable rates for primer extensions, which typically proceeded 5-50%, a DNAP concentration (124 nM) in excess of primer/template (2 nM) had to be employed; thus, the kinetic findings can be viewed as representing virtual single turnover reactions. (We note that DNAP concentrations in excess of primer/template concentration has been used in every kinetic study with Y-Family DNAPs that we are aware of.) Pseudo-first order rate constants are reported (Table 2). Dpo4 was included in every experiment to insure that findings between experiments were comparable. Km was previously shown to be 6.5-25-fold greater than the dNTP concentration (100μM) used herein (see Table 1 in reference ^*(40)*^); thus, rate constants are in a range of the Michaelis-Menten curve where V_max_/K_m_ dominates. BP-adducts are considered stable, but do hydrolyze slowly to give the corresponding BP-tetraol along with the regeneration of guanine in DNA.^*(68)*^ An experiment in Supplementary Materials describes how we established that adduct hydrolysis is not quantitatively significant.

## RESULTS

### Kinetics of dNTP Insertion

Using a template containing -BP-dG, ^32^P-primer extension reactions with dNTPs were performed with purified wt-Dpo4 and wt-Dbh, along with modified mutant derivatives whose amino acid sequences are given in Table 1. Products were separated on DNA sequencing gels and subjected to phosphorimaging.^*(40)*^ Figure 2 shows that Dpo4 inserts all dNTPs significantly opposite –BP-dG, as expected for η/V-class-members, while Dbh preferentially inserts dCTP, as expected for κ/IV-class-members. Figure 2 also shows representative phosphorimages for the other DNAP derivatives.

Phosphorimages were scanned, and Figure 3 shows a representative example. Relative amounts of extended-primer versus unextended-primer were determined from such scans using a method we established,^*(40)*^ and as summarized in Supplemental Materials); in particular, scanning is necessary for accuracy, as other methods (e.g., drawing boxes around bands in phosphoriamges) are highly inaccurate, as we showed.^*(40)*^.

**FIGURE 3:**
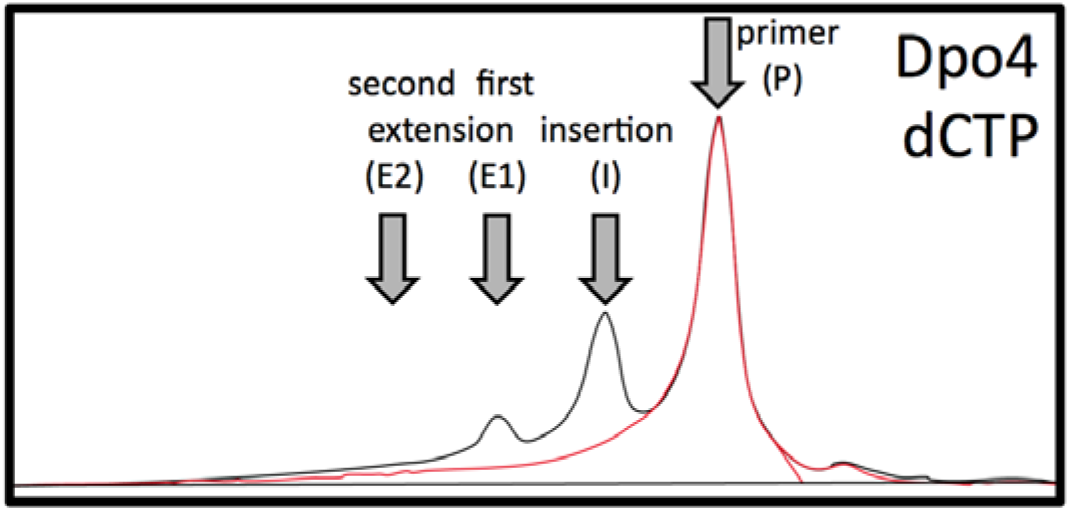
Scan of the dCTP primer extension reaction by SsDpo4 shown in Figure 2. Primer alone is shown in red. The method for computing the fraction of extended primer versus unextended primer is described in detail in reference ^*(40)*^.

A single [dNTP] concentration (100μM) was used, both because many mutant proteins were studied and because all experiments were performed at least in triplicate. Because concentrations of both [dNTP] and [DNAP] (124nM) were in excess of [primer/template] concentration (2nM), reactions followed pseudo-first order kinetics, as we established experimentally.^*(40)*^

For each DNAP/dNTP combination, Table 2 shows the pseudo-first order rate constant (all values × 10^−3^min^-1^) and fractional standard deviation, which is the actual standard deviation divided by the rate constant. The average of all fractional standard deviations is 0.20 (i.e., 20%.)

Because each reaction used the same DNAP and dNTP concentration, comparisons of observed rate constants reveal how protein modifications affect transition state structure/energy.^*(64)*^

### dGTP Misinsertion Has –BP-dG Adduct Bulk in the Major Groove

Y-Family DNAPs have a loop above their active site (the “Active Site Loop” or AS-loop), which for Dpo4 is shown pictorially in Figure 4A and structurally in Figure 4B. The AS-loop for Dpo4 starts at V30 on the minor groove side of the active site (yellow amino acids in Figure 4B), makes a turn and returns to V43 on the major groove side (brown amino acids in Figure 4B). In the active site loop, A42 in Dpo4 (Figure 4B) is aligned with V40 in DNAP IV (Figure 4C), whose R-groups protrude into the major groove just above the templating base. Compared to wt-Dpo4, the mutant A42V-Dpo4 has increased bulk on its major groove side and its dGTP misinsertion (~7.4 × 10^−3^ min^-1^) was suppressed by ~10-fold compared to wt-Dpo4 (~74 × 10^−3^ min^-1^), suggesting that –BP adduct bulk is in the major groove during dGTP misinsertion. The A42V-Dpo4 modification had minimal effect on dCTP, dATP and dTTP incorporations (Table 2), suggesting –BP-dG bulk is not in the major groove in these cases.

**FIGURE 4:**
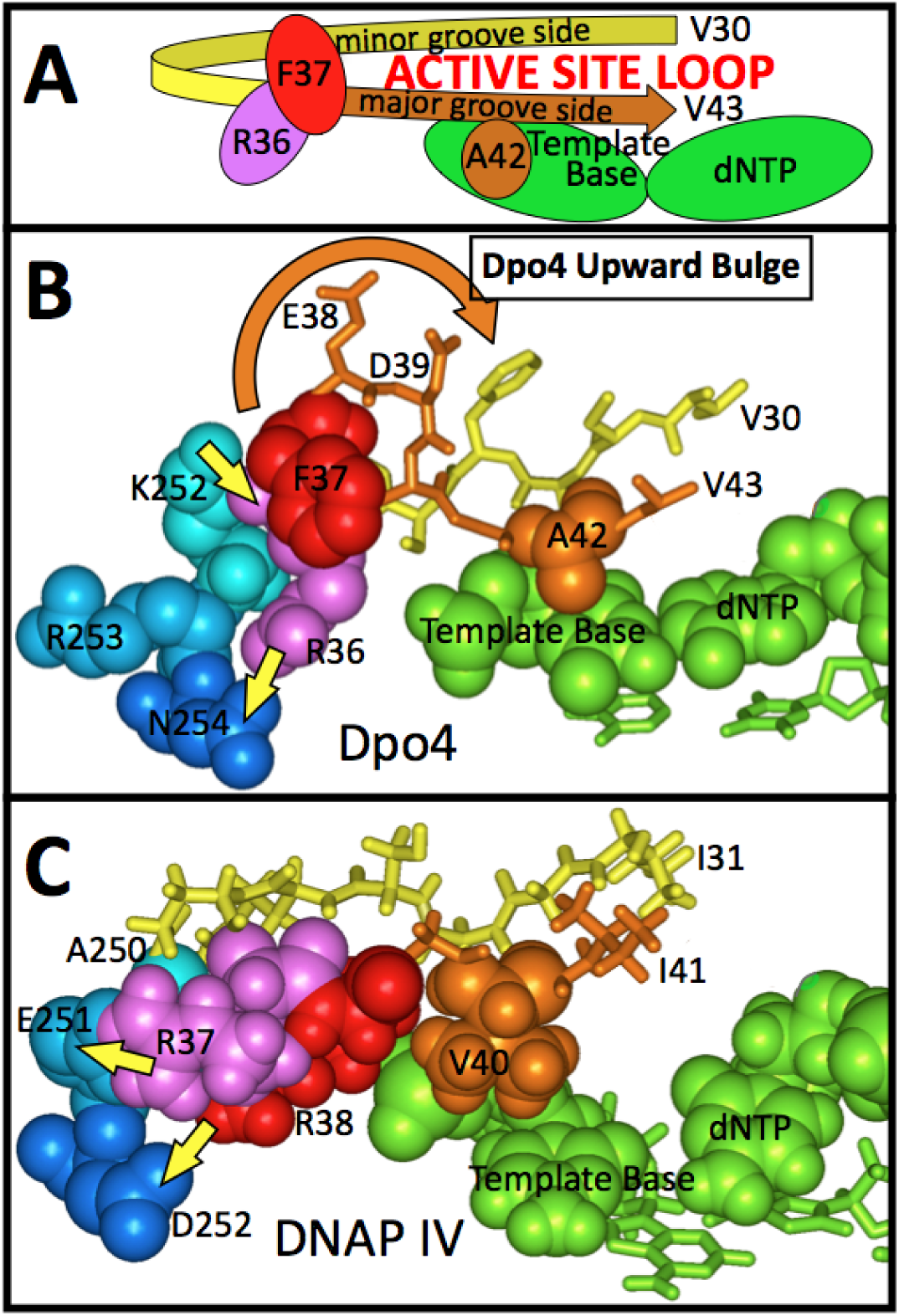
The active site loop (AS-loop) of Dpo4 (V30-V43) is shown pictorially in Panel A and structurally in Panel B, while DNAP IV AS-loop (I31-I40) is shown structurally in Panel C. The view is from the major groove side of the active site. AS-loops in Y-Family DNAPs begin on the minor groove side of the active site, which is represented by amino acids in yellow, and then turns back to the major groove side of the active site, which is represented by amino acids in brown. On the major groove side above the templating base, Dpo4 has A42, while DNAP IV has V40. Dpo4’s AS-NCB between the CC-Domain and LF-Domain-Doman is anchored by two Coulombic interactions: R36/N254 and K252/R36(C=O) (yellow arrows). The AS-loop in DNAP IV has three fewer amino acids than in Dpo4, which are accommodated as a bulge (F37/E38/D39, brown arrow) that moves bulk away from the active site. R36 in Dpo4 (purple) is aligned with R37 in DNAP IV (purple), and the bulge in Dpo4 insures that the next amino acid (F37, red) is above R36 and away from the active site, whereas the next amino acid in DNAP IV (R38, red) is below R37 and closer to the active site. DNAP IV’s AS-NCB between the CC-Domain and LF-Domain-Doman is anchored by two Coulombic interactions: R37/E251 and R38/D252 (yellow arrows).

The AS-loop in Dpo4 (V30-V43) has three more amino acids than in DNAP IV (I31-I41), which are accommodated in Dpo4 as a small upward bulge (Figure 4B, brown arrow) that moves protein mass away from the templating base on the major groove side (as explained in the Figure 4 legend). The three amino acids in this bulge (F37, E38 and D39) were removed to give Δ3-Dpo4, whose dGTP misinsertion velocity (~4.8 × 10^−3^ min^-1^) was ~15-fold slower than wt-Dpo4 (~74 × 10^−3^ min^-1^), while dATP, dCTP and dTTP incorporation rates changed minimally for wt-Dpo4 versus Δ3-Dpo4. Once again, these findings suggest that –BP adduct bulk is in the major groove during dGTP misinsertion, but not during dATP, dCTP and dTTP incorporations.

The entire wt-Dpo4 AS-loop was replaced by the smaller AS-loop from DNAP IV (I31-I41), which lacks the bulge present in Dpo4, to give the mutant IVloop-Dpo4, whose dGTP misinsertion decreased ~11-fold, suggesting again that –BP-dG bulk is in the major groove. In this case, however, dCTP, dATP and dTTP incorporations were affected by addition of the DNAP IV AS-loop to Dpo4. This behavior was investigated previously^*(65)*^ and is discussed below in a subsequent subsection that describes the active site Non-Covalent Bridge (NCB).

The entire wt-Dpo4 AS-loop was replaced by the smaller AS-loop from UmuC (V29-V37), which is the polymerase subunit of EcDNAP V, to give the derivative Vloop-Dpo4, whose dGTP misinsertion rate decreased ~9.5-fold compared to wt-Dpo4. dATP, dCTP and dTTP rates were minimally affected. The smaller UmuC AS-loop is also not expected to have a bulge like in Dpo4.

At its N-terminus, hDNAP κ has an extra 100aa, which forms the U-shaped “N-clasp” (Figure 1C, scarlet ribbon) that restricts access to the major groove side of the active site. ^*(13)*^ hDNAP κ’s N-clasp inhibits insertions opposite adducts with bulk on the major groove side of the active site, ^*(49, 69)*^ and molecular modeling studies have shown why. ^*(70, 71)*^ When the N-clasp was added to wt-Dpo4 to give N-clasp-Dpo4, dGTP misinsertion (~8.1 × 10^−3^ min^-1^) was ~9.1-fold lower than for wt-Dpo4 itself (74 × 10^−3^ min^-1^). Though we have no direct structural evidence that the N-clasp adopted its proper structure in the N-clasp-Dpo4 derivative, this suppression of dGTP insertion is consistent with expectations if the N-clasp is indeed blocking the major groove as it does in hDNAP κ and is, thus, preventing –BP-adduct bulk from being in the major groove during dGTP misinsertion. Furthermore, N-clasp addition had little effect on dCTP insertion (~5.1 × 10^−3^ min^-1^ vs. ~7.2 × 10^−3^ min^-1^), dATP misinsertion (~11.2 × 10^−3^ min^-1^ vs. ~12.2 × 10^−3^ min^-1^) or dTTP misinsertion (~3.8 × 10^−3^ min^-1^ vs. ~3.8 × 10^−3^ min^-1^), which is consistent with the expectation that the N-clasp is indeed blocking the major groove, and –BP-adduct bulk is not in the major groove during dATP, dCTP or dTTP incorporation.

The five findings presented in this subsection are all consistent with the notion that –BP-adduct bulk is in the major groove during dGTP misinsertion, but not during dATP, dCTP or dTTP incorporation.

### dCTP, dATP and dTTP Incorporation Rates Depend on the Size of an Opening on the Minor Groove Side of the Active Site

Canonical base pairing between –BP-N^2^-dG and dCTP requires the BP-moiety to be in the minor groove, since the N^2^-atom is in the minor groove in a Watson-Crick-like base pair. Molecular models revealed that EcDNAP IV has a large opening on the minor groove side of the active site, which helps accommodate -BP-bulk when the adduct-dG moiety is in the active site (Figure 1D)^*(66)*^ and might facilitate dCTP insertion. Recent X-ray structures have shown this large minor groove opening in EcDNAP IV. ^*(17, 26)*^ Furthermore, hDNAP κ has a large slot in the minor groove, which can accommodate BP-bulk.^*(28)*^

In DNAP IV, the lack of R-group bulk on G33 and G74 results in its minor groove opening being large (Figure S1). In the equivalent positions, Dpo4 has V32 and M76, which plug the minor groove opening (Figure S1). V32 and M76 in wt-Dpo4 were converted to glycines to give the double mutant V32G/M76G-Dpo4, and dCTP insertion increased by ~2.4-fold compared to wt-Dpo4. Rates also increased for dATP (~3.5-fold) and dTTP (~3.0-fold) (Table 2). dGTP misinsertion changed minimally (~1.3-fold decrease). This suggests that –BP-bulk is in the minor groove during incorporation of dCTP, dATP and dTTP, but not dGTP.

The AS-loop in hDNAP κ is significantly smaller (G131-L136) than in Dpo4 (V30-V43), DNAP IV (I31-I41) or UmuC (V29-V37), and G131 adopts φ/θ-angles unique to glycines. ^*(72)*^ Both of these features contribute to hDNAP κ having a marge minor groove opening; in fact, it is a slot,^*(13, 28)*^ which is key for accurate bypass of +BP-dG.^*(25, 28)*^ Dpo4’s AS-loop was replaced by hDNAP κ’s AS-loop to give κloop-Dpo4, whose velocities were ~4.2-fold faster than wt-Dpo4 for dCTP and ~3.4-fold faster for dTTP, while dGTP misinsertion is minimally affected. These findings also suggest that –BP-dG has bulk in the minor groove during dCTP and dTTP, but not dGTP incorporation. However, one finding is inconsistent with naïve expectations: dATP misinsertion did not increase with κloop-Dpo4 (11.0x 10^−3^ min^-1^ versus 11.2 × 10^−3^ min^-1^), which suggests either (i) that some feature of the κloop is able to suppress the reaction rate with whatever minor groove –BP-dG conformation is associated with dATP misinsertion, or (ii) that – BP-dG is not in the minor groove. If κloop addition to Dpo4 re-routed dATP misinsertion to a mechanism involving –BP-bulk in the major groove, then addition of N-clasp should suppress dATP misinsertion, which was not observed: the value for N-clasp/κloop-Dpo4 (13.0x 10^−3^ min^-1^) was similar to κloop-Dpo4 (11.0 × 10^−3^ min^-1^). This suggests that possibility (i) is more likely, which is addressed further below.

dGTP misinsertion was similar for N-clasp-Dpo4 (8.1 × 10^−3^ min^-1^) and the double mutant N-clasp/κloop-Dpo4 (7.6 × 10^−3^ min^-1^). Similarly, dATP misinsertion was similar for N-clasp-Dpo4 (8.1 × 10^−3^ min^-1^) and the N-clasp/κloop-Dpo4 (7.6 × 10^−3^ min^-1^). dCTP insertion increased somewhat for N-clasp/κloop-Dpo4 (from 7.2 × 10^−3^ min^-1^ to 13.1 × 10^−3^ min^-1^), implying that κloop addition improved fidelity for dATP and dGTP misinsertion slightly. However, dTTP misinsertion increased ~3.1-fold (from 3.8 × 10^−3^ min^-1^ to 12.0 × 10^−3^ min^-1^), making this fidelity improvement purine-specific.

### A Non-Covalent Bridge Near the Active Site Between the LF-Domain and the CC-Domain Suppresses dATP/dGTP/dTTP Misinsertions

An Active Site Non-Covalent Bridge (AS-NCB) in Dpo4 (Figure 1A) includes two Coulombic interactions: (1) The guanidinium moiety of R36 in the AS-loop interacts with N254 in the LF-Domain, and (2) the backbone carbonyl-oxygen of R36 interacts with the ammonium moiety of K252 in the LF-Domain (Figure 4B, yellow arrows).

The significance of the AS-NCB to dNTP misinsertion was probed by eliminating both Coulombic interactions: the double mutant R36A/K252A-Dpo4 had higher activity compared to Dpo4 for all three misinsertions: dATP (~4.5-fold), dGTP (~3.7-fold), and dTTP (~2.7-fold), while dCTP changed minimally compared to wt-Dpo4. These findings show that the AS-NCB helps suppress dATP, dGTP and dTTP misinsertion, while minimally affecting correct dCTP insertion.

Considering the single mutants, no dNTP rate was significantly affected by K252A-Dpo4 compared to wt-Dpo4, while with R36A-Dpo4, dATP increased (~3.3-fold), while dCTP, dGTP and dTTP rates changed minimally. This finding is considered in greater depth in Discussion.

The AS-loop and AS-NCB is much different in Dpo4 vs. DNAP IV.^*(65)*^ For DNAP IV, we showed that R37 interacts with E251 and R38 interacts with D252 (Figure 4C), with the major consequence being that DNAP IV’s AS-NCB suppresses dATP, dGTP and dTTP misinsertions more effectively than the AS-NCB of Dpo4 by ~5-fold (see reference 65).

### Distal to the Active Site a Non-Covalent Bridge (Distil-NCB) Between the LF-Domain and the CC-Domain Suppresses dATP/dGTP/dTTP Misinsertions

Dpo4 is an η/V-class-member, while Dbh is a κ/IV-class-member, based on insertion patterns opposite lesions, including a thymine dimer and an AP site. ^*(20, 27, 73)*^ All four dNTPs are incorporated more slowly by Dbh than Dpo4, though misinsertions are slowed more significantly for dATP (~18-fold), dGTP (~67-fold) and dTTP (~7.8-fold) compared to dCTP (~2.6-fold). Thus, Dbh can also be classified a κ/IV-class-member based on insertion patterns opposite –BP-dG.

Studies have shown that a chimera with the LF-Domain of Dbh joined to the CC-Domain of Dpo4 (Dbh/LF-Domain-Dpo4) behaved more like Dbh than Dpo4 with a thymine dimer and an AP site. ^*(20, 27, 73)*^ The same tendency is observed with –BP-dG: Dbh/LF-Domain-Dpo4 misinsertion decreased with dATP (~8-fold), dGTP (~34-fold) and dTTP (~7-fold) compared to wt-Dpo4. dCTP insertion decreased also, but less so (~2.5-fold).

To probe what in Dbh’s LF-Domain might confer κ/IV-class properties on Dpo4, Dpo4 and Dbh were compared, and the greatest difference in amino acid sequence in the LF-Domain is:

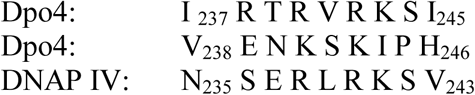

These nine amino acids from Dbh were placed into Dpo4 to give aa238-246/Dbh-Dpo4, which decreased misinsertion in comparison to wt-Dpo4 for dATP (~2.2-fold), dGTP (~7.2-fold) and dTTP misinsertion (~1.9-fold), while dCTP insertion was minimally affected. While our work was in progress, another group showed that a Dpo4 derivative containing approximately this same region behaved more like Dbh with respect to misinsertion on undamaged DNA and insertion opposite an AP site. ^*(20)*^

In X-ray structures ^*(20, 27, 73)*^, Dbh forms a non-covalent bridge (NCB) between the LF-Domain and the CC-Domain on the minor groove side ~8Å distal to the active site (Figure 1B, blue arrow). One key contact is V101/I244—the latter amino acid being in this region. This Distal-NCB is likely to be the structural element that can suppress dATP, dGTP and dTTP misinsertions opposite –BP-dG.

Our DNAP IV molecular model was built via homology modeling using a Dpo4 starting structure. ^*(44, 72)*^ Dpo4 has no Distal-NCB (Figure 1A), and yet a Distal-NCB always formed in our DNAP IV models during molecular dynamics. DNAP IV’s Distal-NCB formed because P99 kinks a β-strand in the CC-Domain, forcing this β-strand ~3Å closer to the LF-Domain compared to Dpo4, thus bringing R240 in DNAP IV’s LF-Domain close enough to interact with residues in DNAP IV’s CC-Domain. At the equivalent position Dpo4 also has arginine (R242), so we investigated whether we had correctly surmised the importance of P99 in DNAP IV by studying the double mutant I101P/A102L-Dpo4, whose rates (compared to wt-Dpo4) decreased for dATP (~2.8-fold) and dGTP (~3.5-fold). (In DNAP IV, L100 fills a hydrophobic pocket to help anchor P99, which is why the double mutant was studied.) dCTP and dTTP insertions were minimally affected. Thus, the addition of I101P/A102L to Dpo4 also suppressed two of three dNTP misinsertions (though not as effectively as the aa238-244/Dbh change) suggesting that a DNAP IV-like Distal-NCB can also improve fidelity in Dpo4.

## DISCUSSION

### Four Different -BP-dG Adduct Conformations Appear to Be Responsible for the Four Different dNTP Insertions

Dpo4 incorporates all dNTPs opposite –BP-dG at significant rates (Figure 2, Table 2). One mechanism for this could be random insertion; i.e., –BP-dG binds in a single non-informational conformation, which Dpo4 cannot interpret, so it merely incorporates whatever dNTP happens to be in its active site. This mechanism is unlikely, given the following observations, which suggest that each dNTP insertion mechanism involves a unique adduct conformation.

[1]dGTP misinsertion has –BP-dG bulk in the major groove, as it was suppressed by protein structural elements that decreased the size of the major groove opening near the active site (e.g., conversion of A42 to valine), while its velocity was unaffected by changes on the minor groove side.

[2]In contrast, dCTP insertion has –BP-dG bulk in the minor groove, as it was affected by protein modifications on the minor groove side (e.g., adding V32G/M76G to Dpo4), but not the major groove side of the active site.

[3]dATP and dTTP misinsertion also involved –BP-dG bulk in the minor groove, because their rates can be affected by protein modifications on the minor groove side, but not on the major groove side. However, the –BP-dG minor groove conformations cannot be the same for dATP and dTTP compared to dCTP, because the presence of an AS-NCB and/or a Distal-NCB between the LF-Domain and the CC-Domain suppressed dATP and dTTP misinsertions, but not dCTP insertion.

[4]Protein modifications to Dpo4 tended to affect dATP and dTTP misinsertion rates similarly, suggesting that their –BP-dG conformations are more similar; however, two findings distinguished dATP versus dTTP misinsertion. [i] The rate for dTTP misinsertion was greater for κloop-Dpo4 than for wt-Dpo4, while dATP misinsertion was similar for κloop-Dpo4 and wt-Dpo4. [ii] dTTP misinsertion only increased when both the R36/N254 and the K252/R36(C=O) interactions involving the AS-NCB were removed; in contrast removal of the R36/N254 interaction alone significantly suppressed dATP misinsertion. These two differences in dATP versus dTTP misinsertion pattern undoubtedly reflect some difference in –BP-dG conformational positioning in the minor groove, given that both dATP and dTTP must sit similarly in the active site.

We proposed^*(74)*^ that different mutations arising from a single BP-dG adduct were caused by different BP-dG adduct conformations, as reviewed extensively.^*(75)*^ The analysis in this subsection, suggesting that the four different dNTP insertion pathways arise from four different – BP-adduct conformations, provides further support for this hypothesis. The nature of these four conformations vis-à-vis the four dNTP insertions will be addressed elsewhere.

#### Additional Analysis of the Major Groove Opening

Adding protein bulk to the opening on the major groove side of Dpo4 decreased dGTP misinsertion, via the A42V modification, via the removal of the three amino acid bulge in Dpo4’s AS-loop, or addition of the N-clasp addition.

Others have noted that the N-clasp in DNAP κ inhibits bypass of adducts with bulk on the major groove side, ^*(49, 69)*^ and molecular modeling studies have shown why. ^*(70, 71)*^ Still others have speculated that having bulk at the position equivalent to A42 in Dpo4 might sterically exclude certain kinds of lesions; e.g., M135 in DNAP κ was proposed to help exclude the entry of thymine dimers into the active site from the major groove side. ^*(16)*^ Our finding—that the small bulge in Dpo4’s AS-loop moves protein bulk away from the templating base—has not been noted previously and is a structural feature present in other η/V-class-members (discussed below).

#### Non-Covalent Bridges Orient the LF-Domain to Prevent Misinsertion Opposite Aberrant Adduct Structures

The AS-NCB and Distal-NCB are on the minor groove side of the active site, so it is not surprising that they suppress dATP and dTTP misinsertions, which have –BP-dG bulk in the minor groove. However, the AS-NCB and Distal-NCB do not suppress dCTP insertion, even though it also has –BP-dG bulk in the minor groove. Furthermore, the AS-NCB and Distal-NCB <u>*do*</u> suppress dGTP misinsertion, which has bulk in the major groove.

Thus, the formation of non-covalent bridges on the minor groove side per se cannot explain their selective ability to suppress all three misinsertions. It is more likely that these NCBs help position the LF-Domain so it can exclude aberrant DNA structures from entering the active site, independent of whether the aberrant structures have adduct bulk in the minor groove or the major groove. Others have shown that the Distal-NCB is likely to be important for Dbh functioning vis-à-vis other bypass events. ^*(20, 27, 73)*^

##### Generalizations about Y-Family DNAP Structure

Dpo4 has an upward bulge on the major groove side of its active site (Figure 4C: F37/E38/D39); our findings suggest that this bulge allows adduct-bulk to be better accommodated on the major groove side of the active site based on our findings for dGTP misinsertion opposite -BP-dG. While this bulge promotes dGTP misincorporations opposite –BP-dG, which is counter-productive, it appears to be helpful for allowing entry of other lesions (e.g.) UV-damage) into the active site. We note that human and yeast DNAP η have upward bulges analogous to the one in Dpo4, ^*(16, 21)*^ suggesting that this feature might be important in many η/V-class-members.

η/V-class Y-Family DNAPs have plugged minor groove openings, as observed with Dpo4 (V32 and M76 in Figure S1). The minor groove openings are even more plugged in human-and yeast-DNAP η, which have, respectively, unique seven and thirty-four amino acid inserts that completely cap their respective minor groove openings. ^*(16, 21)*^ The minor groove opening is also plugged with UmuC.^*(72)*^

By simply grafting the N-clasp from hDNAP κ onto Dpo4, we generated a derivative (N-clasp-Dpo4) that suppressed dGTP misinsertion, thus improving fidelity compared to wt-Dpo4 itself (Table 2). We were not optimistic that this experiment would work as envisioned, though molecular modeling suggested that it might (data not shown); furthermore, we do not even know if these extra 100 amino acids are truly operating in N-clasp-Dpo4 analogously to how they operate in hDNAP κ. Nevertheless—by whatever mechanism—the findings show that if a Dpo4-like DNAP were simply to acquire an N-clasp-like sequence from some genetic source at some point during evolution, then fidelity of BP-adduct bypass would improve. Thereafter, one can speculate that the AS-loop might have shrunk—this supposition being based on the observation that κloop addition to N-clasp-Dpo4 (giving κloop/N-clasp-Dpo4) improved both dATP and dGTP fidelity. Next, DNAP κ’s N-clasp might have acquired its non-covalent bridge (Figure 1C, scarlet arrow), which is on the major groove side of the active site, along with the losses of the minor groove side. While this pathway for the generation of DNAP κ’s basic structure is obviously highly speculative, it seems sensible.

In summary, κ/IV-class-members have greater protein bulk on the major groove side of their active sites and more open space on the minor groove side, while η/V-class-members are the opposite. To achieve these goals, prokaryotic and Archaea Y-Family DNAPs principally vary the number of amino acids and/or R-Group bulk in the AS-Loop. Eukaryotic Y-Family DNAPs have also accomplished these goals, though via the grafting on of significant protein inserts; e.g., the N-clasp to block the major groove in DNAP κ, and the aforementioned inserts in DNAP η (seven and thirty-four amino acids, respectively, for yeast and humans) to block the minor groove opening.

## ACKNOWLEDGEMENTS

We thank Dr. Nicholas Geacintov for providing the –BP-dG-containing oligonucleotide, and Dr. Roger Woodgate for supplying several Dpo4-and Dbh-containing plasmids, along with Drs. Louise and Satya Prakash for supplying a plasmid containing the gene for human DNAP κ.

## SUPPORTING INFORMATION

Supplemental information is provided.

## AUTHOR INFORMATION

### Notes

The authors declare no competing financial interests.

## ABBREVIATIONS

^2^BP: benzo[*a*]pyrene
+BP-dG: [+ta]-BP-N^2^-dG (Figure 1)
-BP-dG: [-ta]-BP-N^2^-dG (Figure 1)
AP site: apurinic/apyrimidinic site
thymine dimer: a cyclobutanepyrimidine dimer formed between adjacent thymimes by UV light
DNAP: DNA polymerase
CC-Domain: catalytic core domain that includes the thumb/palm/fingers domains, which operates as a unit and contains the active site
LF-Domain: the little finger domain, which helps grip DNA
SsDpo4: *Sulfolobus Solfataricus* Dpo4
SaDbh: *Sulfolobus acidocaldarius* Dbh
EcDNAP IV: *Escherichia coli* DNA polymerase IV
AS-NCB: active site non-covalent bridge
Distal-NCB: Distal non-covalent bridge

